# Ascorbate concentration in *Arabidopsis thaliana* and expression of ascorbate related genes using RNAseq in response to light and the diurnal cycle

**DOI:** 10.1101/138008

**Authors:** William Laing, Cara Norling, Di Brewster, Michele Wright, Sean Bulley

## Abstract

We explore where transcriptional regulation of ascorbate concentration lies in plants. Is it in biosynthesis,recycling, regulation or consumption? Arabidopsis thaliana plants were grown under controlled environment at four photon flux density levels (PFD). Rosettes from plants were harvested at the four PFD levels and over a diurnal cycle and after a step change in PFD and analysed for ascorbate concentration and transcript levels measured by RNAseq. Ascorbate concentrations and expression of genes in the L-galactose ascorbate biosynthesis, recycling,consumption pathways and regulation are presented to provide a full analysis of the control of ascorbate by environmentally modulated gene expression. Ascorbate concentration responded to PFD levels but not to time of day and showed only a small response to change of PFD after 2 days. Of the L-galactose pathway genes, only GDP galactose phosphorylase (GGP) showed a significant response in to different PFDs, time of day and to change in PFD. Other genes also showed limited responses. This study compares gene expression of a range of ascorbate related genes to changes in environment in a unified way and supports the concept that GGP is the key regulatory gene in ascorbate biosynthesis and that post transcriptional regulation is also important.

**Highlight:** In a comprehensive study of expression of all ascorbate related genes the data is consistent with the control of leaf ascorbate concentration by transcription being through the expression of GDP galactose phosphorylase.

## Introduction

Ascorbate or Vitamin C is an important redox agent in plants and animals, an enzyme co-substrate, and an essential nutrient (vitamin) for humans and some other animals. The regulation of ascorbate concentration in plants is complex with transcription, translation, protein turnover and other factors suggested (Bulley and Laing, 2016a). Control would influence biosynthesis, recycling, transport and consumption of ascorbate. However while ascorbate concentration is normally well regulated within a tissue or organ ofa species, it varies strongly between different species and with environmental conditions, and in some plants, concentrations are very high (Bulley and Laing, 2016b), suggesting that their concentrations may not be always well constrained by these factors.

It has been established that ascorbate concentration and some genes implicated in its biosynthesis respond to light and temperature and time of day (Gatzek *et al.,* 2002; Tamaoki *et al.,* 2003; Muller-Moule *et al.,* 2004; Bartoli *et al.,* 2006; Dowdle *et al.,* 2007; Yabuta *et al.,* 2007; Maruta *et al.,* 2008; Müller-Moulé, 2008; Kotchoni *et al.,* 2009; Page *et al.,* 2012; Conklin *et al.,* 2013; Heyneke *et al.,* 2013; Wang *et al.,* 2013; Noshi *et al.,* 2016a) (Gao *et al.,* 2011) as well as development (Ioannidi *et al.,* 2009; Bulley *et al.,* 2009b; Li *et al.,* 2010). In general, ascorbate concentration and gene expression increase with increased photon flux density (PFD) and show some increase from lights on until later in the day. Significant changes also occur during fruit development (Bulley *et al.,* 2009b). However, the published data tends to be limited in scope, for example, only covering selected genes in response to environment or during fruit development, both using PCR to measure expression.

Previous work by us using transformation, gene expression or mapping has established that http://submit.biorxiv.org/control of ascorbate concentration resides at least partially with the enzymes GDP galactose phosphorylase (GGP) and in a synergistic fashion with GDP mannose epimerase (GME) (Laing *et al.,* 2007; Bulley *et al.,* 2009a; Bulley *et al.,* 2012; Mellidou *et al.,* 2012; Laing *et al.,* 2015) which has been supported by others (Zhou *et al.,* 2012; Li *et al.,* 2013) (Gilbert *et al.,* 2009; Massot *et al.,* 2012) (Yoshimura *et al.,* 2014). However other L-galactose pathway biosynthetic enzymes have also been suggested to control ascorbate concentration including galactose-1-P phosphatase in tomato (GPP) (Ioannidi *et al.,* 2009) and GDP mannose pyrophosphorylase in acerola (Badejo *et al.,* 2007). Myo-inositol oxygenase, an enzyme in an alternative pathway, has also been proposed to strongly stimulate ascorbate in *Arabidopsis* (Lorence *et al.,* 2004). Recycling of oxidised ascorbate has also been implicated in determining ascorbate concentration (Chen *et al.,* 2003; Naqvi *et al.,* 2009) (Gest *et al.,* 2013). If these enzymes are critical in determining ascorbate concentration it is likely regulatory control will be at least partially at the transcriptional level.

We undertook a comprehensive targeted approach to evaluate the control of ascorbate concentrations in leaves of *Arabidopsis thaliana* by gene expression with a focus on the biosynthesis, regulation, recycling and consumption of ascorbate. We grew and harvested plants in environmental conditions and at times that were expected to affect ascorbate concentration through biosynthesis, recycling and/or degradation. We included both high light stresses at low night temperature as well as a change in light intensity, and undertook a diurnal series of harvests. We used RNAseq to measure the total gene expression in *Arabidopsis* leaves. We then selected previously identified genes relating to ascorbate biosynthesis and metabolism, including the genes that encode the enzymes of the L-galactose biosynthesis pathway, genes that encode for recycling enzymes, genes that encode for enzymes that consume ascorbate and genes that encode regulators (Bulley and Laing, 2016a). From this global data we calculated the expression level of these genes. For biosynthetic genes we restricted ourselves to the L-galactose pathway of ascorbate biosynthesis as at least in *Arabidopsis* this is the main and critical pathway of ascorbate biosynthesis (Dowdle *et al.,* 2007; Lim *et al.,* 2016).

This study examines leaf gene expression of a range of ascorbate related genes in relation to ascorbate concentration under changes in environment and time of day in a unified way and supports the hypothesis that expression of genes for especially GDP-L-galactose phosphorylase *(GGP)* and somewhat GDP mannose epimerase *(GME)* are the central to the regulation of ascorbate biosynthesis. We also show that ascorbate concentration is well regulated in that its concentration does not vary rapidly under short term perturbations in gene expression (including time of day and change in PFD) but is well acclimated to the long term environment. Our data also supports the post transcriptional regulation of ascorbate concentration.

## Materials and Methods

### Plant growth

*A. thaliana* ‘Columbia’ plants were grown for 6 weeks in two controlled environment rooms at the New Zealand Controlled Environment Laboratory (Warrington and Mitchell, 1975), Palmerston North. Prior to sowing, seed were stratified at 5°C for three days in water. The seed were sown by hand pipette into 70 mm diameter pots containing a coarse Daltons™ peat bark potting mix, top layered with fine Daltons™ potting mix (bark/peat/pumice). The pots were base soaked in water and left to germinate for 3 days in the dark at a constant 23°C. Following germination, seedlings were overhead misted until their true leaves emerged. Then the pots were thinned to one plant per pot before the six leaf stage, giving a total of 432 plants per room. Plants were fed nutrients by base flooding, then allowed to drain, using half strength modified Hoagland’s solution (Brooking, 1976) and grown at 23 ±0.5°C at 70% relative humidity (0.8 kPa water vapour deficit (VPD)), approximately 380 ppm CO2, 252 μmoles photons m^-2^ s^-1^ at a 10 h day length. Light was provided by a mixture of two 1 kW high intensity discharge (‘Metalarc’, GTE; Sylvania, UK) and four 1kW tungsten halogen (Thorn, Enfield, UK) lamps.

After 4 weeks, the plants were uniform, with a ~50 mm diameter rosette and 13 leaves. At this stage, environmental conditions were adjusted to stress the plants. The PFD was increased in the rooms to 700 μmoles photons m^-2^ s^-1^ for 10 h photoperiod using a mixture of six 1kW high intensity discharge lamps and six 1kW tungsten halogen lamps in both rooms, and four levels of light using three levels of neutral shade cloth were established (plus unshaded) providing PFDs at plant height of 100, 250, 450 and 700 μmoles m^-2^ s^-1^. One room remained constant at 23°C night and day, the other at a diurnal of 23°C day and 5°C night. Other conditions were the same as for the establishment period, except VPD was 0.3 kPa for the low night temperature room. Plants at the two highest PFD levels at the low night temperatures developed a noticeable red tinge, due to anthocyanin (Page *et al.,* 2012) to their leaves. No flowering was observed in plants due to the short days.

After 2 weeks of growth under these conditions, destructive harvests were made over the next 48 h as described in the results. Plants were harvested at one time point over the four PFD levels (PFD treatment); plants grown at 250 μmoles m^-2^ s^-1^ were harvested over two diurnal cycles at approximately 3 h intervals (diurnal treatment); and plants were also subjected to a step change in PFD from 100 to 700 for approximately 20 h (step treatment). Depending on the treatment between four (PFD transfer and some diurnal measurements) and six (PFD response and some diurnal measurements) separate plants were harvested as replicates and processed separately for ascorbate and RNA extraction. For ascorbate measurements the plants were analysed separately, for RNAseq, RNA samples were pooled to create two biological replicates after the separate RNA extraction. Harvested rosettes were rapidly wrapped in aluminium foil and immersed in liquid nitrogen to freeze samples. Leaf samples were powdered under liquid nitrogen and were transferred to storage at- 80°C until further processing.

### Ascorbate measurements

Ascorbate was measured by extracting *ca.* 100 mg samples of powdered material into 0.8% w/v metaphosphoric acid, 2 mM EDTA and 2 mM Tris(2-carboxyethyl)phosphine hydrochloride (TCEP HCL) and quantifying ascorbate by HPLC. The samples were centrifuged at 14,000g for 15 minutes to clarify the extract and then analysed by HPLC using a rocket column (Altima C18 3 micron from Phenomenex (Auckland New Zealand) at 35 C. Ascorbate was quantified by injecting 10 μL into a Dionex Ultimate® 3000 Rapid Separation LC system (Thermo Scientifc). Instrument control and data analysis was performed using Chromeleon v7.2 (Thermo Scientific). Solvent A was 5 mL methanol, 1mL 0.2M EDTA pH 8.0 and 0.8mL o-phosphoric acid in 2 L. Solvent B was 100% acetonitrile. The flow as 1.0 mL/min and the linear gradient started with 100% A and B was increased to 30% at 4.5 min, then to 90% B at 6 min. The column was then washed with 100% B and then returned to 100% A. The column was monitored at 245 nm and ascorbate quantified by use of authentic standards.Ascorbate was verified by its UV spectrum. This method gave the sum of oxidized and reduced ascorbate, namely total ascorbate. as described earlier (Laing *et al.,* 2015). Each plant was measured separately giving 4 to 6 biological replicates per treatment.

### RNA extraction and processing

Approximately 100 mg of ground plant material were extracted using the Sigma-Aldrich Spectrum Plant Total RNA kit using the protocol described by Sigma-Aldrich. RNA was then treated with DNAse RNA quality was assessed using a Nanodrop 1000 spectrophotometer (Thermofisher Scientific) as well as a 2100 Bioanalyzer system (Agilent), using samples with a RNA integrity number > 7. Samples within a time period and treatment were randomly combined to create two replicates per treatment measured. Because of financial constraints, only samples from the low night temperature room were processed and selected time points from the diurnal series giving an even spread of times. Paired-end Illumina RNA-seq sequencing (125 bp length), including library preparation, of the 24 separate RNA samples was undertaken by New Zealand Genomics Ltd (NZGL: Otago, New Zealand). Samples were run over two lanes to avoid lane effects. Both the forward and reverse direction paired end sequencing was undertaken, giving 96 separate files in total.

### Bioinformatics analysis

Data was processed using the Galaxy pipeline (Goecks *et al.,* 2010). First sequences were trimmed (14 bp removed from the 5’ end, 5 from the 3’ end leaving about 105 bp). Data were converted to the Sanger format using FASTQ Groomer, filtered by quality and Bowtie2 run to align reads to Arabidopsis mRNA data from TAIR 10. The status of the alignment file checked by Flagstat. Over 92 to 95% of reads were of acceptable quality. The protocol SAM/BAM to counts was then applied to count the number of aligned sequences that matched selected genes in the Arabidopsis TAIR10 database for a particular experiment (e.g. the response to PFD). Data for a particular set of genes (e.g. the Galactose pathway of ascorbate biosynthesis) was then selected from this count data and output for further analysis. The paired end reads and duplicate lane sequencing data was combined to create an average and this was regarded as a single biological replicate to be analysed with the separate biological replicate (i.e. n = 2).

Statistical analysis was carried out using the PredictMeans package in R (https://cran.r-project.org/web/packages/predictmeans/predictmeans.pdf) to calculate LSD between means or using a general linear model in MINITAB to calculate standard errors and the significance of differences using the Tukey method. Both methods gave the same estimates of significance and only the MINITAB results are presented.

Raw and processed RNAseq data for all Arabidopsis genes has been deposited with NCBI GEO under accession number GSE94995.

## Results

### Physiological and biochemical measurements

#### Plant Growth

*Plant Growth*. was significantly reduced by the lower night temperature and increased somewhat with increased PFD, especially at the 23°C night temperature (Fig. 1). The interaction between PFD and night temperature was highly significant (P < 0.001), as was the direct effect of PFD and night temperature (P < 0.001).

**Figure 1.**
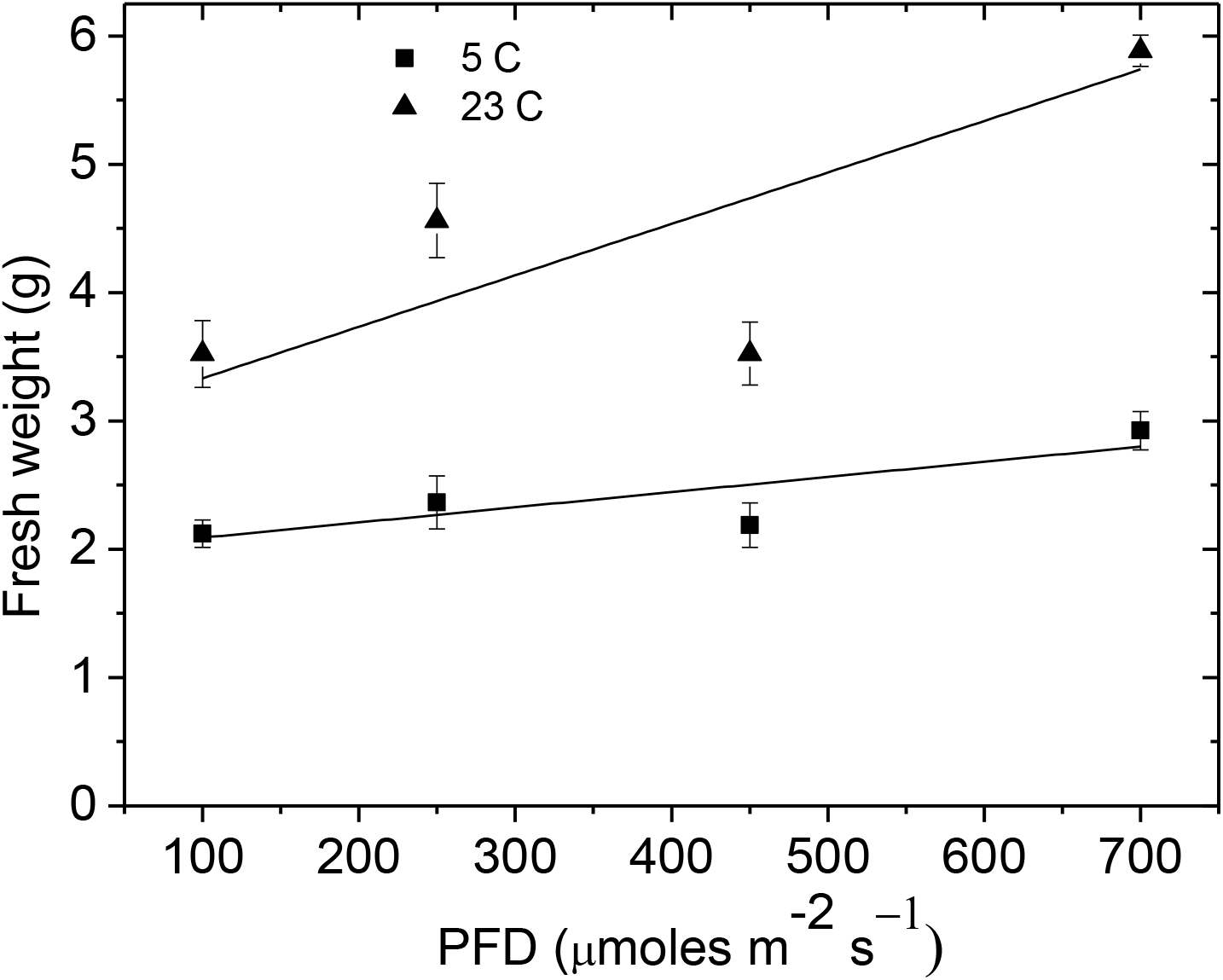
Plant growth as measured by rosette fresh weigh at different growth Photon flux density (PFD) levelsand night temperatures. Triangles represent weights of plants grown at 23/23°C and squares are plants grown at 23/5°C.

#### Ascorbate

Ascorbate was measured in rosettes from six week old plants grown for 2 weeks at 23/23°C or at 23/5°C and a range of PFD levels. Ascorbate increased with PFD, especially at the lower night temperature (Fig. 2A; Table 1), and the interaction between growth PFD and night temperature was significant (P< 0.001).

**Figure 2.**
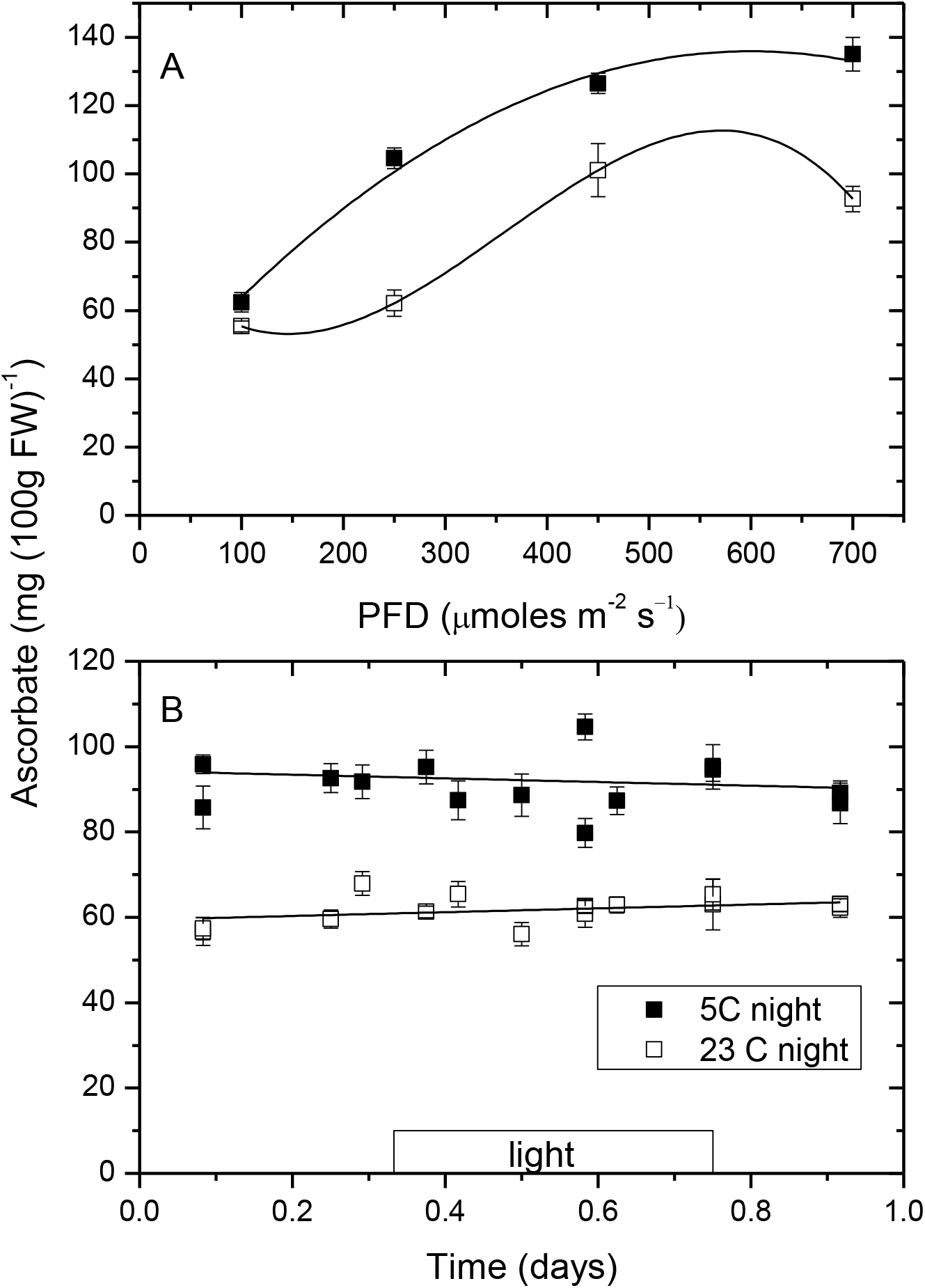
Response of leaf ascorbate concentration to PFD (A; See also Table1) and to time of day for plants grown at 250 μmol m^-2^ s^-1^ (B). In graph A,leaves were sampled at 0.58 days, approximately half way through the light period. In graph B, plants were sampled over three days but the data is plotted over one day. Bars represent standard errors of the mean. Lines in A are second (5°C night) and third order (23°C night) polynomial fits to the data. Lines in B are linear fits to the data.

**Table 1.** Statistical analysis of the response of ascorbate concentration to Photon Flux Density (PFD) and night growth temperature.

| PFD $\mu\text{mol m}^{-2} \text{s}^{-1}$ | Night Temp. ($^{\circ}\text{C}$) | N | Mean<br>mg/100g FW | Grouping<br>($P < 0.05$) |
| --- | --- | --- | --- | --- |
| 100 | 5 | 6 | 62.4 | C |
| 250 | 5 | 6 | 104.6 | B |
| 450 | 5 | 6 | 126.5 | A |
| 700 | 5 | 6 | 135.1 | A |
| 100 | 23 | 7 | 55.3 | C |
| 250 | 23 | 7 | 62.7 | C |
| 450 | 23 | 6 | 101.1 | B |
| 700 | 23 | 6 | 92.7 | B |

Night temperature and PFD and their interaction were all significant (P < 0.001). Means that do not share a letter are significantly different using Tukey method and 95% confidence using a general linear model withfactors PFD, night temperature and their interaction

No diurnal trend in ascorbate concentration for plants grown at 250 μmol m^-2^ s^-1^ was seen (Fig. 2B; time ns, night temperature *P* < 0.001). Again, ascorbate was higher for plants grown at a low night temperature.

We also measured the effect of transferring plants from 100 to 700 μmol m^-2^ s^-1^ at both temperatures. Plants were transferred at 0.4 days soon after the lights came on then sampled intensivelythe rest of that day and again at about midday the next day (Fig. 3). Even on the second day, only a small effect of change in light intensity on ascorbate was seen in these transferredplants, with a 19% (5 C) and 28% (23 C) rise in ascorbate compared to the base 100 μmol m^-2^ s^-1^ concentration. This represents only 51% and 72% of the steady state ascorbate concentration for plants grown at 700 μmol m^-2^ s^-1^ respectively, compared to 46% and 60% for plants grown constantly at the two temperatures at 100 and 700 μmol m^-2^ s^-1^. Ascorbate concentration was rising but slowly, similar to previous results (Muller-Moule*et al.,* 2003).

**Figure 3.**
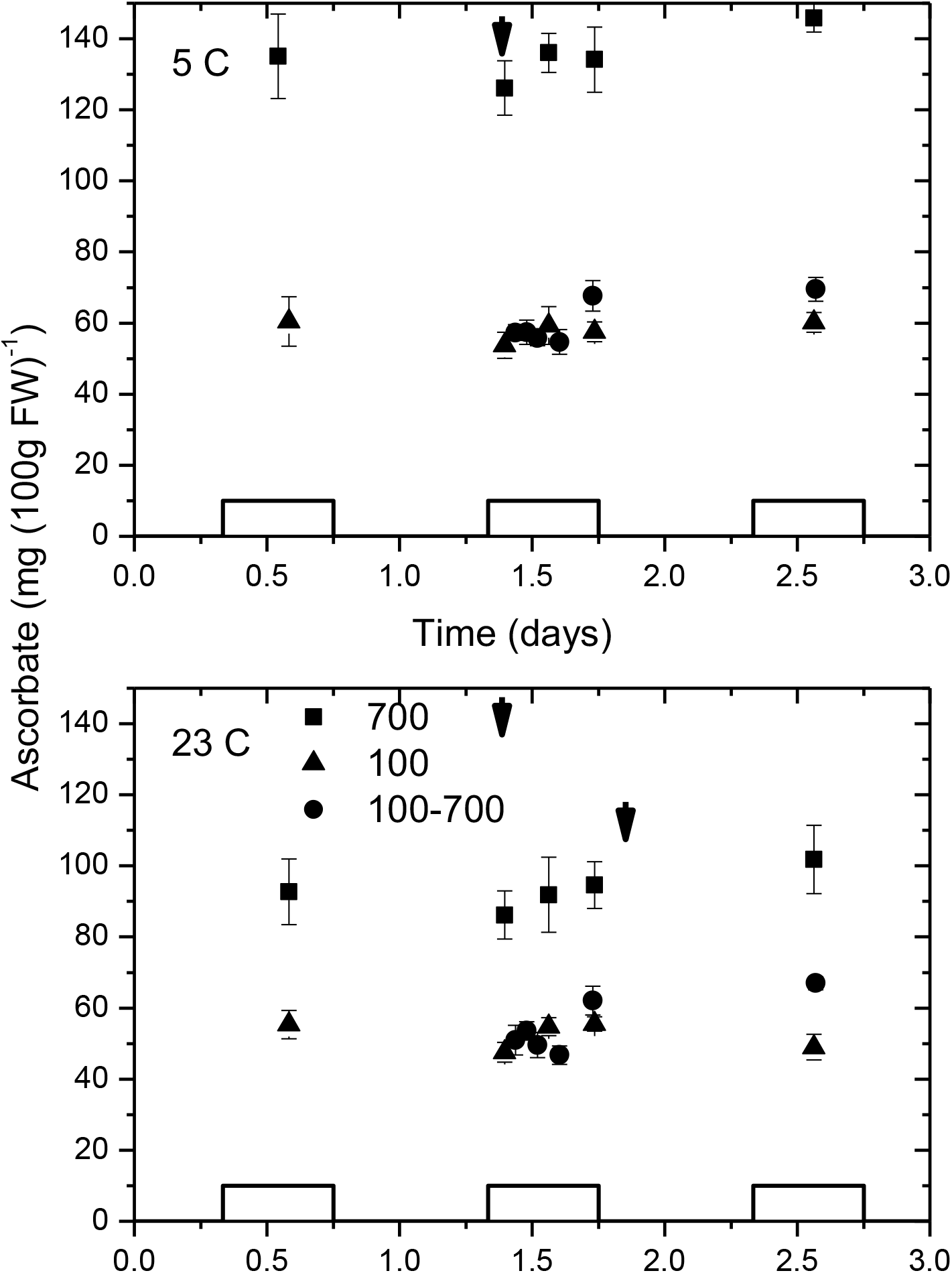
Response of leaf ascorbate to a change in PFD from 100 to 700 μmol m^-2^ s^-1^ for plants grown at 23 and 5°C night temperature. The small black arrow at the top of each graph shows the time when plants were transferred (0.4 days). Closed squares are 700 μmol m^-2^ s^-1^ grown plants, triangles 100 μmol m^-2^ s^-1^ grown plants and circles plants that were transferred from 100 to 700 μmol m^-2^ s^-1^. Bars represent SEM and the 3 boxes on the X axis represent the times when lights were on.

### Gene expression

#### Verification genes

In order to verify the gene expression data, we compared the response of selected genes to the published literature. For example, to validate the circadian rhythm data for our ascorbate related genes, we selected a range of genes where published data showed different circadian responses (Schaffer *et al.,* 2001; McClung, 2006; Harmer, 2009; Hsu and Harmer, 2014; Romanowski and Yanovsky, 2015; Nolte and Staiger, 2015). These included circadian responsive genes and photosynthetic genes. For example, *gigantea (GI)* transcripts accumulate towards the middle of the day (Panigrahi and Mishra, 2015) while genes such as *CCA1* show morning peaks (Nagel *et al.,* 2015). These responses to time of day are shown for five genes in Fig. 4 and show excellent agreement with the literature.

The response of Rubisco small subunit genes to PFD and time of day at 250 μmol m^-2^ s^-1^ was also analysed. The gene expression generally showed a declining expression with increasing growth PFD and little response to time of day, similar to that previously reported (Dean *et al.,* 1989) (Fig. S1). Three photosystem genes were also selected and are plotted in Fig. S2.

**Figure 4.**
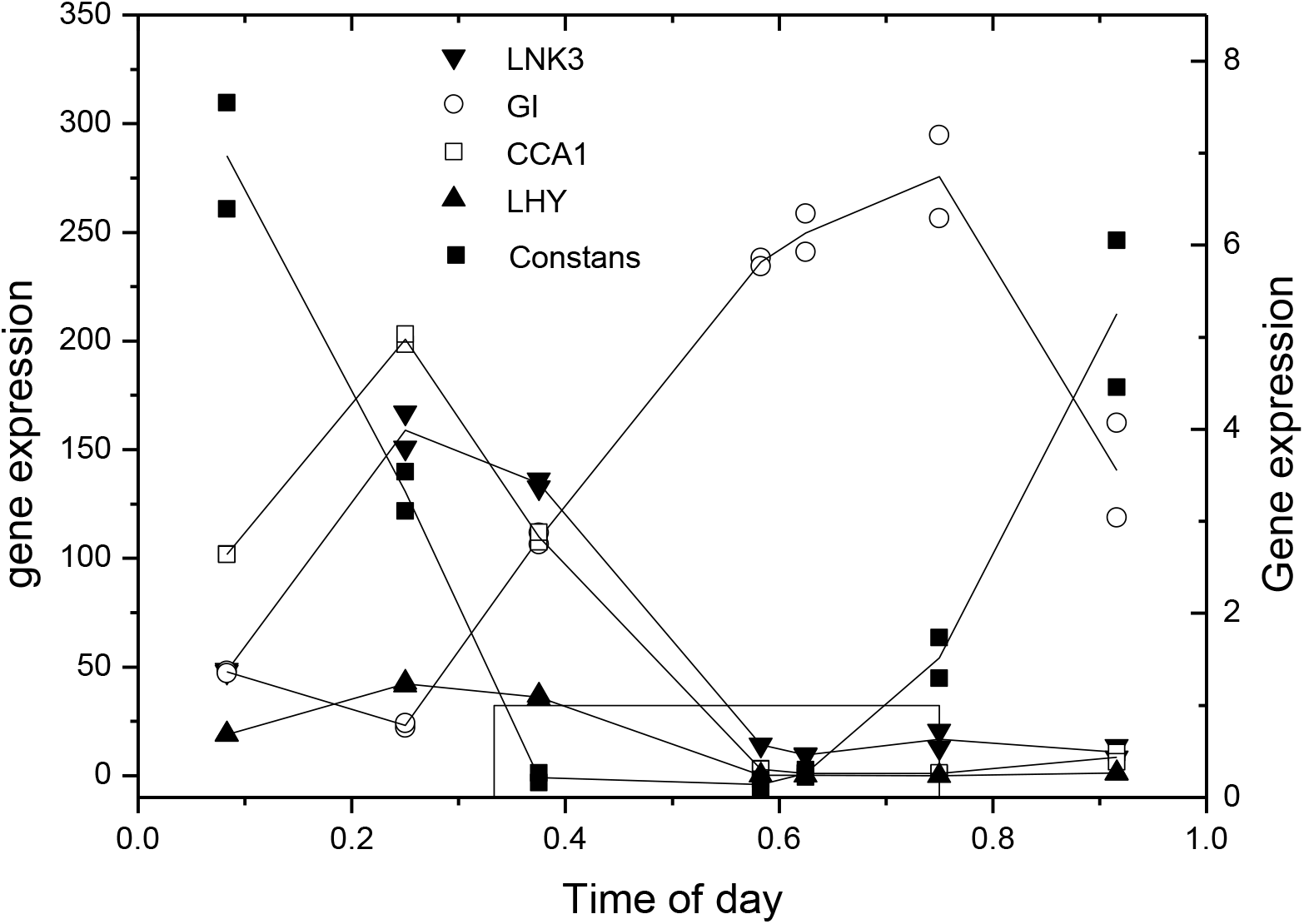
The response of gene expression (Reads Per Kilobase of transcript per Million mapped reads (RPKM)) to time of day for a selection of genes previously shown to have diurnal rhythms in gene expression for plants grown at 250 μmol m^-2^ s^-1^. The genes are LNK3 (AT3G12320) (downward filled triangle),GI (AT1G22770) (open circles), Constans (AT5G15840) (filled squares, RH scale), CCA1 (AT2G46830) (open squares),and LHY/ AT1G01060 (upward closed triangles). Each time point is represented by two individually plotted independent biological replicates. The square box like line at the bottom of the graph represents when the lights were on.

These show a strong diurnal trend in agreement to literature observations. Expression also declines significantly with increasing PFD as did the RubisCO small subunit genes. The decline in photosynthetic gene expression, especially the light harvesting genes (Bailey *et al.,* 2001), with increasing PFD may reflect the high and near saturating PFD the plants were grown under.

The concordance of this expression data with published or expected results supports the reliability of the sequencing data.

### Ascorbate related genes

#### L-galactose biosynthesis genes

Response to growth PFD: Many of the genes in the L-galactose pathway showed no significant change in gene expression between different growth PFDs. These included *phosphoglucoisomerase (PGI), phosphomannose isomerase (PMI), phosphomannomutase (PMM), L-galactose phosphate phosphatase (GPP), galactose dehydrogenase (GDH),* and *galactono lactone dehydrogenase (GLDH)* (Fig. 5A). Previously identified genes for enzymes that have been suggested to regulate the L galactose pathway, namely GDP mannose pyrophosphorylase *(GMP), GME* and *GGP* (VTC2 version) all showed significant response to PFD, with the gene expression rising with PFD (Fig. 5B). The other version of *GGP (GGP2, VTC5),* showed a smaller response to PFD.

**Figure 5.**
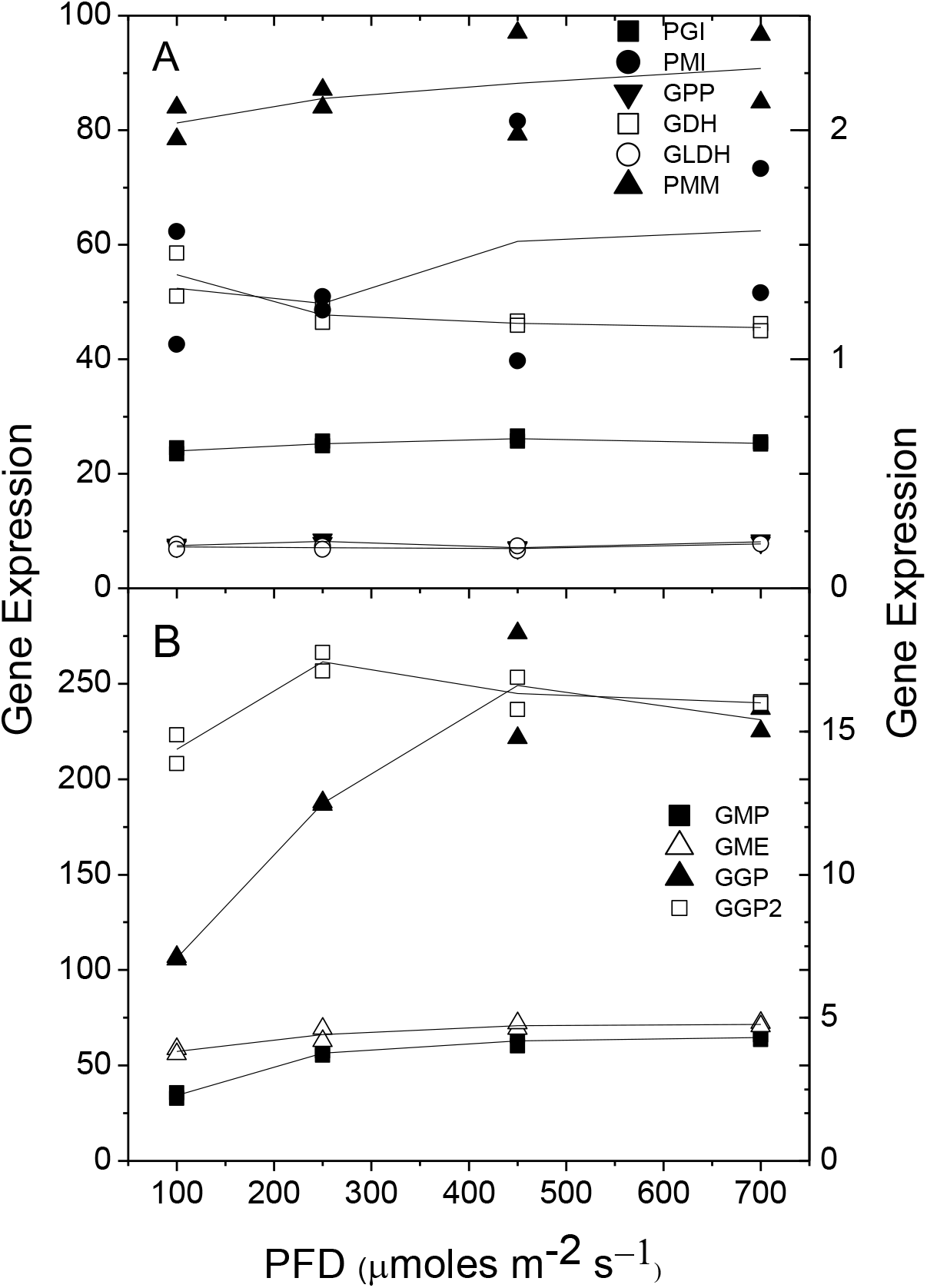
The response of gene expression (RPKM) of ascorbate biosynthetic genes to growth PFD. The two biological replicates are plotted separately and the lines follow the mean of these replicates. Samples were taken at 0.583 days. Graph A. The following non PFD responsive genes are plotted (with the F values from ANOVAR): Phosphoglucoisomerase *(PGI;* filled square. F=0.084); Phosphomannose isomerase *(PMI:* filled circle, right hand axis. F=0.87); phosphomannomutase *(PMM:* filled upward triangle. F=0.685); L-galactose phosphate phosphatase *(GPP:* Downward filled triangle. F=0.052); Galactose dehydrogenase *(GDH:* open square. F=0.094); and galactono lactone dehydrogenase *(GLDH:* open circle. F=0.049). Graph B. The following PFD responsive genes are plotted: GDP mannose pyrophosphorylase *(GMP:* solid square. F<0.001); GDP mannose epimerase *(GME:* open upward triangle. F=0.022); GDP galactose phosphorylase *(GGP:*upward solid triangle. F=0.007); GDP galactose phosphorylase 2 *(GGP2:* open square,righthand axis. F=0.027).

Response to time of day for plants grown 250 μmol m^-2^ s^-1^ (diurnal treatment): Transcription of all the genes in the L-galactose pathway showed response to time of day except *PGI,* but often the response was small *(PMI, PMM, GMP, GGP2, GPP* and *GLDH;* Fig.6A). The most striking diurnal changes were seen with *GGP, GMP, GDH* and *GME* (Fig. 6B).

**Figure 6.**
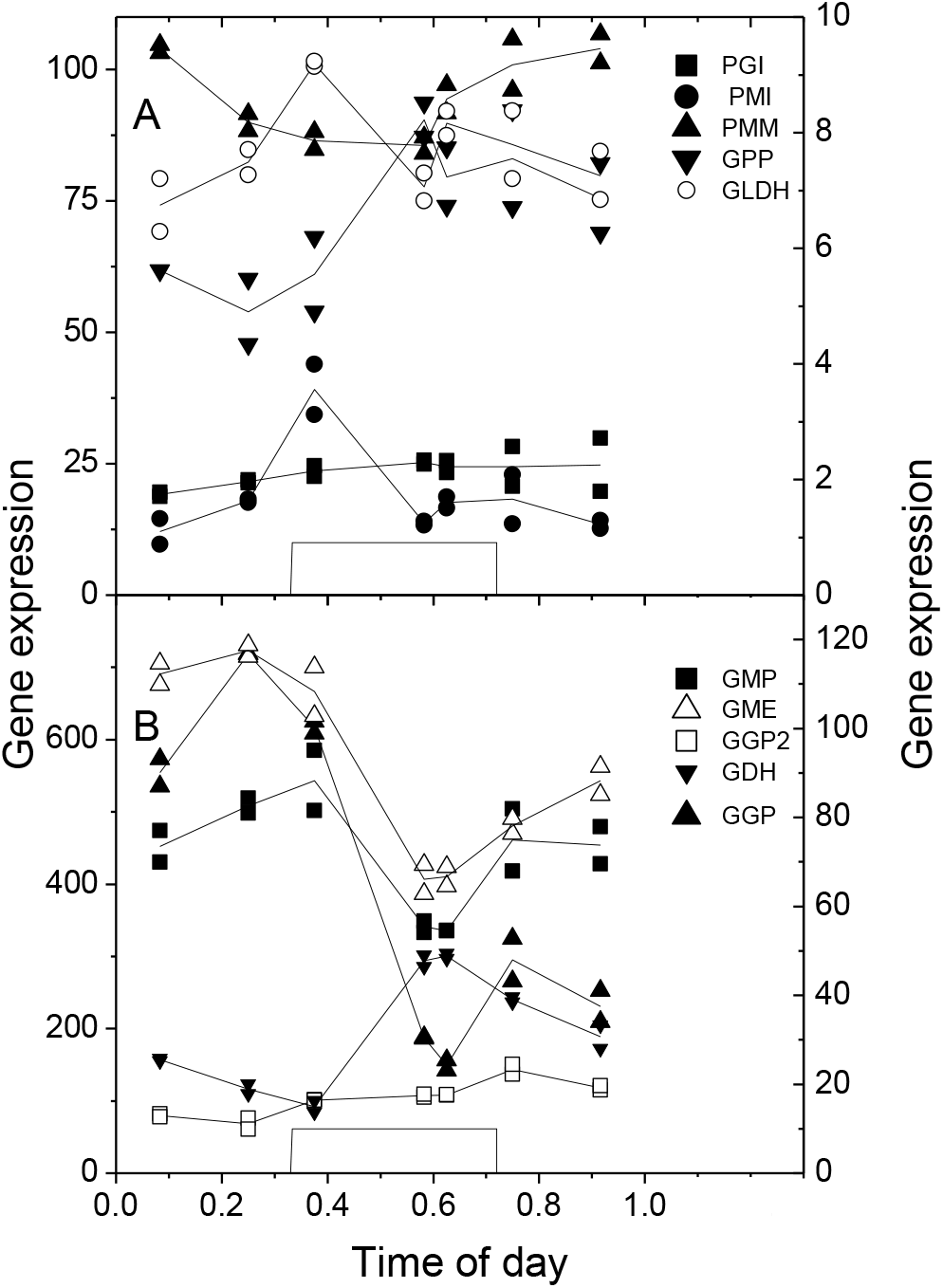
The response of gene expression (RPKM) of ascorbate biosynthetic genes to time of day. The two biological replicates are plotted separately and the lines follow the mean of these replicates. Plants were grown at 250 μmoles photons m^-2^ s^-1^: The square wave shows when the lights were on. Graph A.The following genes that show minimal response to time of day are plotted (F values from ANOVAR in brackets): Phosphoglucoisomerase (*PGI*; filled square. F=0.604); Phosphomannose isomerase *(PMI:* filled circle, right hand axis. F=0.003); phosphomannomutase *(PMM:* filled upward triangle. F=0.004); L-galactose phosphate phosphatase *(GPP:* Downward filled triangle, RH axis. F=0.028); and galactono lactone dehydrogenase *(GLDH:* open circle, RH axis. F=0.023). Graph B. The following genes that responded more strongly to time of day (except GGP2) are plotted: GDP mannose pyrophosphorylase *(GMP:* solid square, RH axis. F=0.006); GDP mannose epimerase *(GME:* open upward triangle, RH axis. F<0.001); GDP galactose phosphorylase *(GGP:* upward solid triangle. F<0.001); GDP galactose phosphorylase 2 *(GGP2:* open square, RH axis. F<0.001); Galactose dehydrogenase *(GDH:* downward triangle. F<0.001).

Interestingly, GDH showed a peak in the middle of the day while the other peaked before the lights came on.

Response to a step change in PFD for plants grown at 100 μmol m^-2^ s^-1^. We measured gene expression after a step change in PFD from 100 to 700 μmol m^-2^ s^-1^. While ascorbate was slow to respond to changed demand due to increased PFD (Fig. 3A), several genes *(GGP, GMP* and *GME*) showed a strong response over the next two days to increased PFD (Fig. 7, Table S1).

**Figure 7.**
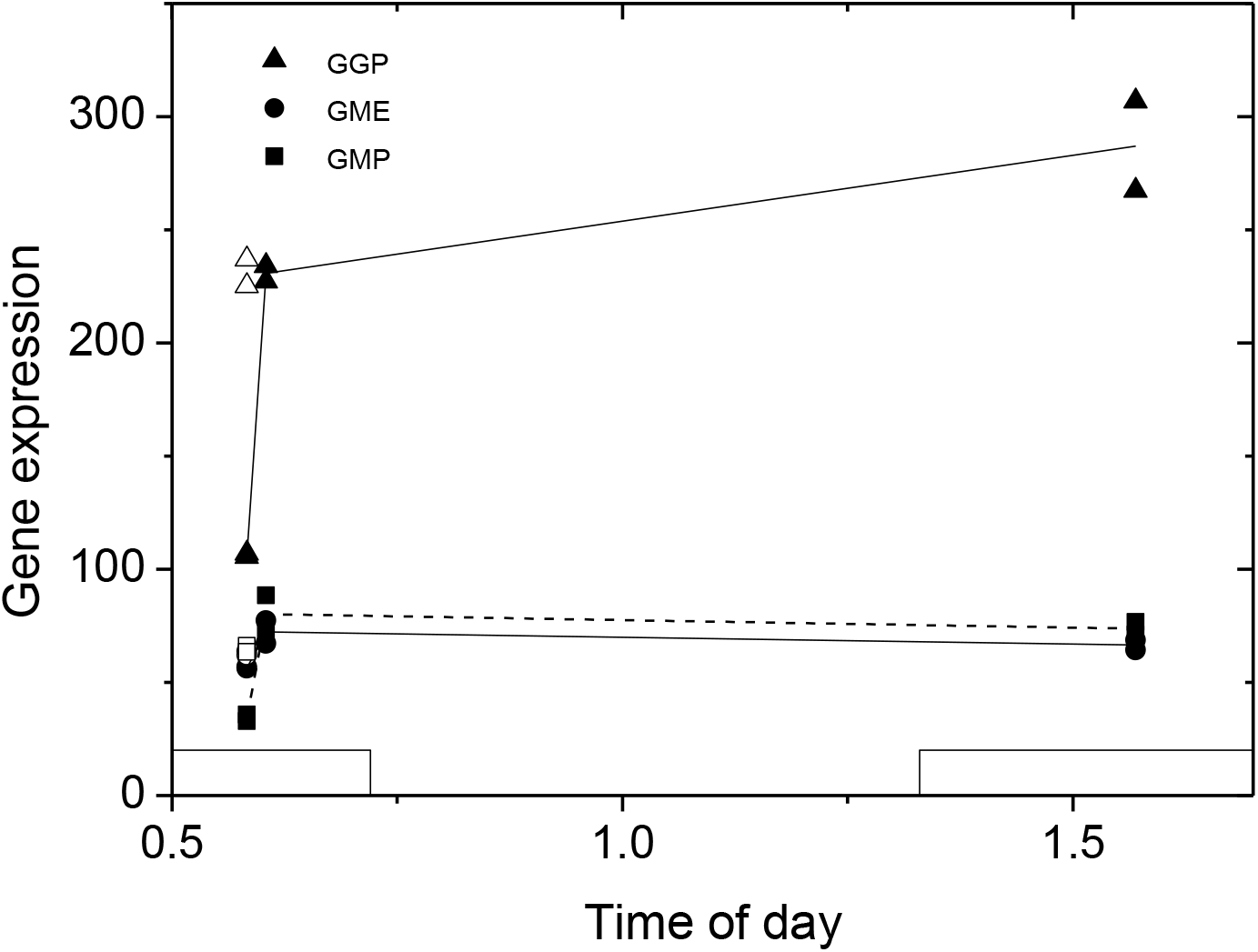
Change in gene expression of ascorbate biosynthetic genes (RPKM) after a step change from 100 to 700μmol m^-2^ s^-1^. The PFD was increased at time = 0.40 and gene expression measured at 1.6 day and at 2.57 days (i.e. approximately 1.2 and 2.2 days later). The initial measurement at 0.58 is for plants grown at 100 μmol m^-2^ s^-1^. The two biological replicates are plotted separately and a line is drawn to join their mean values from the 100 μmol m^-2^ s^-1^starting point. The isolated pairs of open symbols at time = 0.58 are the gene expression for the same genes for plants grown constantly at 700 μmol m^-2^ s^-1^. Squares represent *GGP,* circles *GMP* and triangles *GME.* Full data is shown in Table S1. The square wave on the X axis represents when the lights were on each day.

#### Ascorbate recycling genes

##### Response to growth PFD

We measured the expression of genes related to ascorbate recycling including *dehydroascorbate reductase* (DHAR) genes, *monodehydroascorbate reductase* (MDAR) genes and *glutathione reductase* (GR) genes. In response to growth PFD, some genes within a group showed an increased expression at higher PFD, while others were non responsive (Fig. 8 and Table S2). In the case of *DHAR* genes, the gene of one chloroplast located protein responded strongly to PFD, while another did not, and in the case of *MDAR* genes, both genes of chloroplast located proteins responded to PFD. There was little response of the gene of the chloroplast located *GR* to growth PFD.

##### Response to time of day (diurnal trends)

Neither *DHAR* nor *MDAR* genes showed large changes in expression during the day (Fig. 9 and Table S3). However, GR genes did show a larger decrease in expression during the day, peaking around lights on.

**Figure 9.**
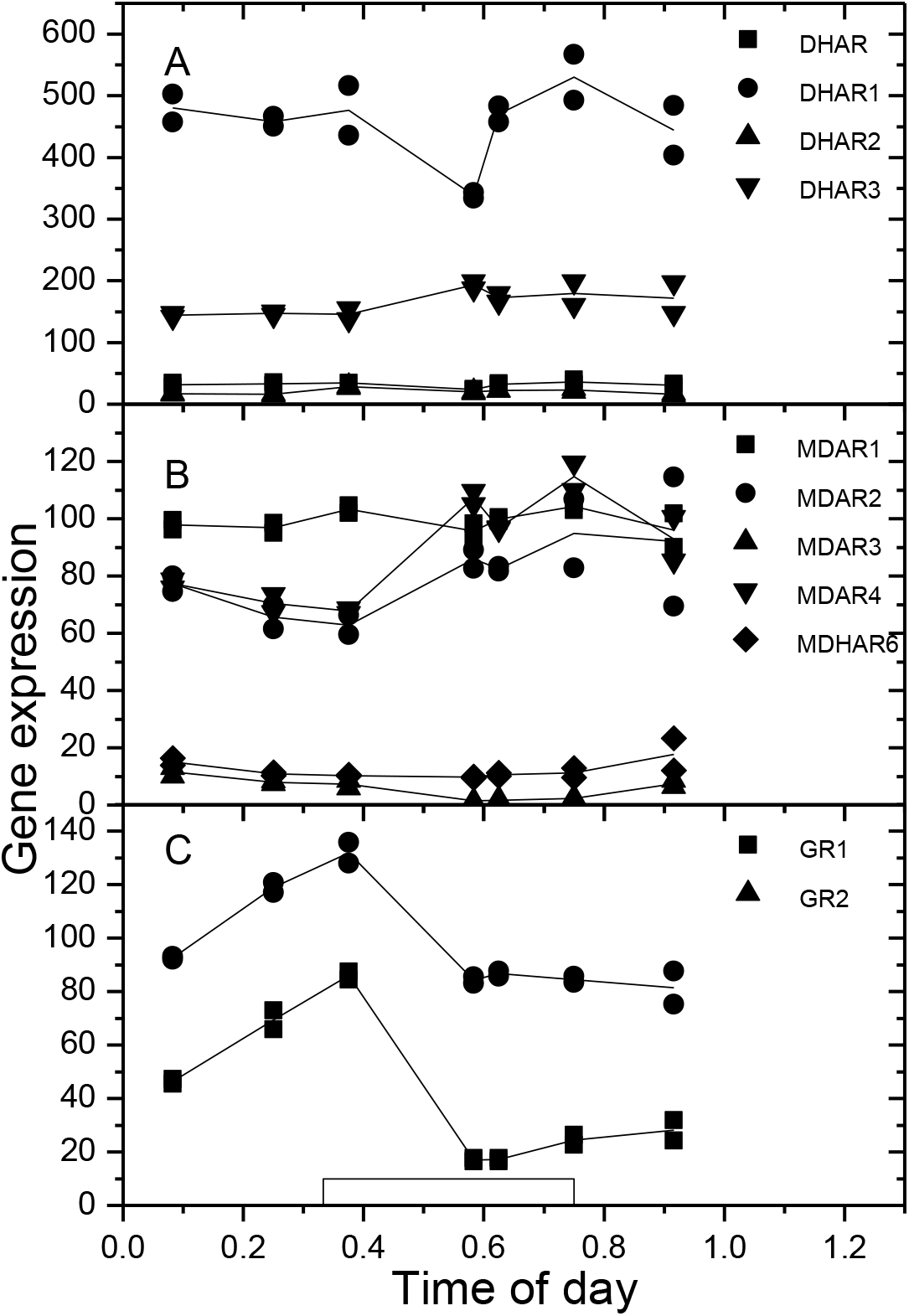
The response of gene expression of ascorbate recycling genes to time of day. The two biological replicates are plotted separately and the lines follow the mean of these replicates. Plants were grown at 250 μmoles photons m^-2^ s^-1^: The square wave in C shows when the lights were on. Graph A, Dehydroascorbate reductase *(DHAR)* genes (squares are *DHAR (AT1G19550),* circles are*DHAR1 (AT1G19570),* upward triangles are *DHAR2 (AT1G75270)* and downward triangles are *DHAR3 (AT5G16710));* Graph B, Monodehydroascorbate reductase *(MDAR)* genes (squares are*MDAR1 (AT3G52880),* circles are*MDAR2 (AT5G03630),* upwards triangles are*MDAR3 (AT3G09940),* downward triangles are*MDAR4 (AT3G27820)* and diamonds are*MDHAR6 (AT1G63940));* Graph C, Glutathione reductase (GR) genes (squares are *GR1 (AT3G24170),* circles are *GR2 (AT3G54660)).* Statistical significances of changes are shown in Table S4.

##### Response to a step change in PFD

Some *DHAR* and *MDAR* genes that were expressed at a higher level in high PFD responded to increased light with increased gene expression (Fig. 10A; Fig. 10B) reflecting the results seen in Fig. 8A and Fig. 8B. The results with the DHAR genes reflect that seen elsewhere on a step change in PFD (Noshi *et al.,* 2016b). The GR genes generally increased after a step change in PFD even though they showed little to no response to PFD (Fig. 10C compared to Fig. 8C).

**Figure 10.**
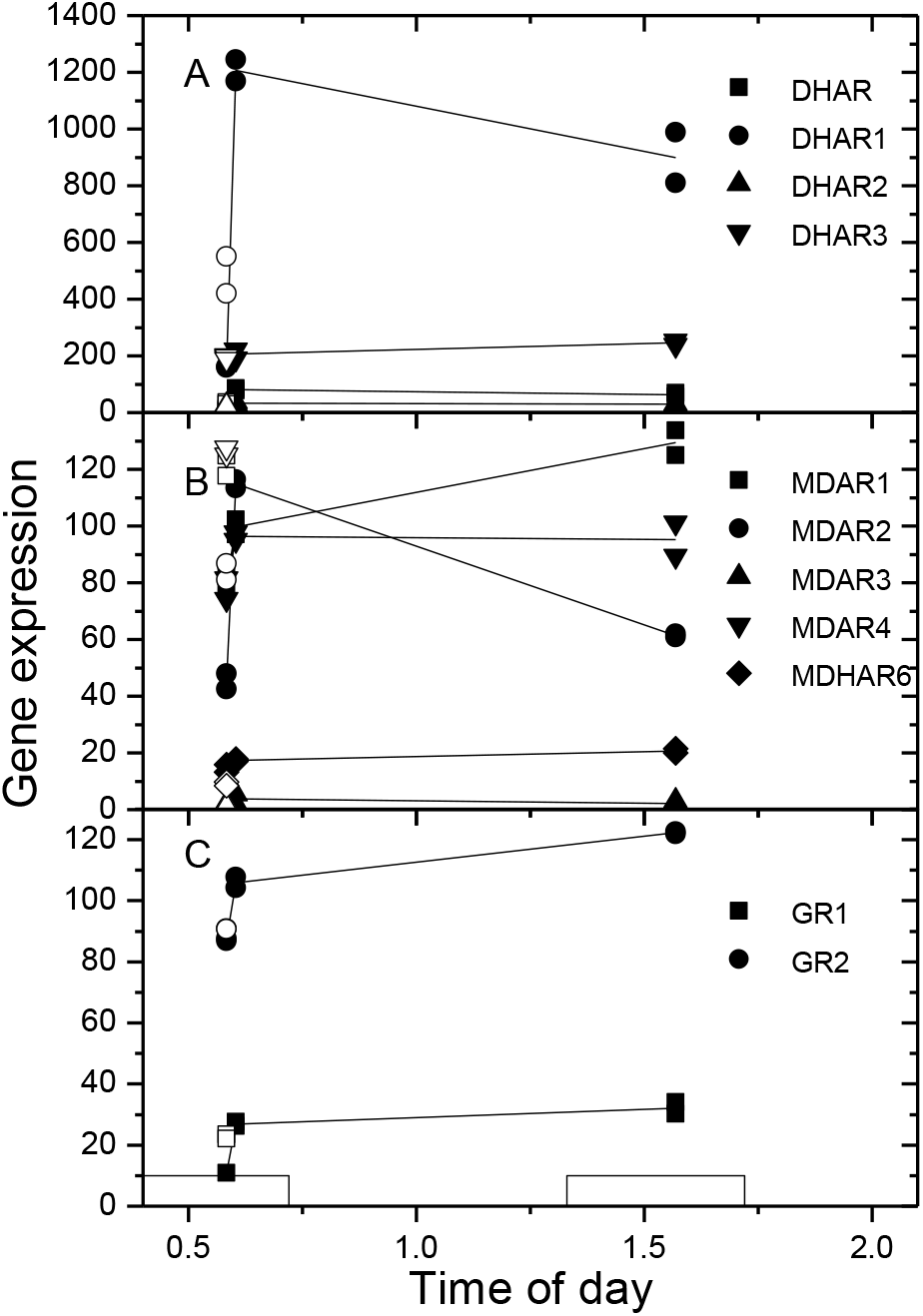
The response of gene expression of ascorbate recycling genes to a step change in PFD. The two biological replicates are plotted separately and the lines follow the mean of these replicates. The PFD was increased at time = 0.40 and gene expression measured at 1.6 day and at 2.57 days (i.e. approximately 1.2 and 2.2 days later). The initial measurement at 0.58 is for plants grown at 100 μmol m^-2^ s^-1^.Open symbols represent steady state values of gene expression in 700 μmol m^-2^ s^-1^grown plants taken at the same time of day as samples from plants that had a step change in PFD. Closed symbols represent plants that had a step change in PFD. Graph A, Dehydroascorbate reductase *(DHAR)* genes(squares are *DHAR (AT1G19550),* circles are *DHAR1 (AT1G19570),* upward trianglesare *DHAR2 (AT1G75270)* and circles are *DHAR3 (AT5G16710);* Graph B, Monodehydroascorbate reductase *(MDAR)* genes (squares are *MDAR1 (AT3G52880),* circles are*MDAR2 (AT5G03630),* open triangles are *MDAR3 (AT3G09940),* downward triangles are *MDAR4 (AT3G27820)* and diamonds are *MDHAR6 (AT1G63940));* Graph C, Glutathione reductase (GR) genes (squares are *GR1 (AT3G24170),* triangles are *GR2 (AT3G54660)).* Statistical significances of changes are shown in Table S5.

**Figure 8.**
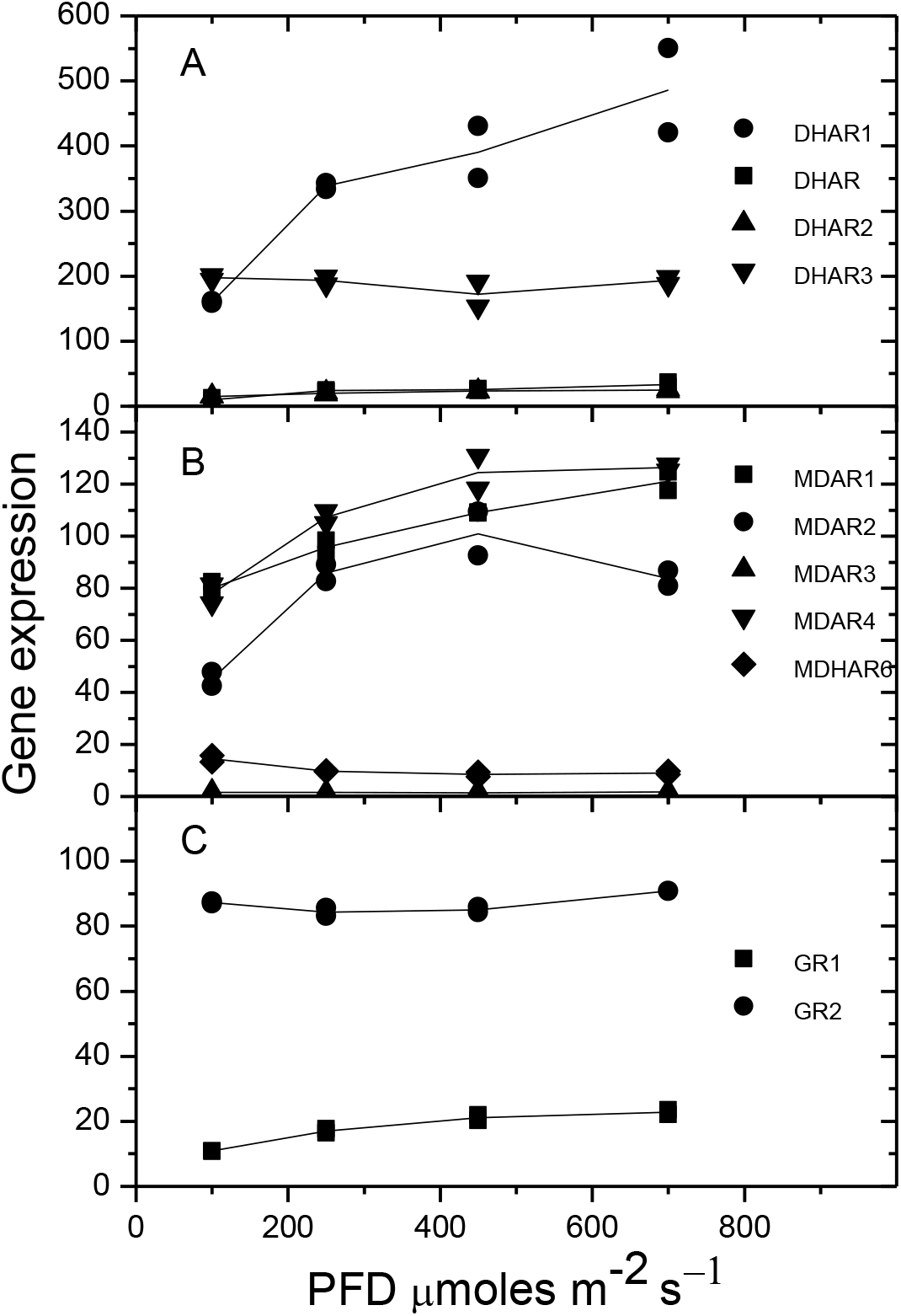
The response of gene expression (RPKM) of ascorbate recycling genes to growth PFD. The two biological replicates are plotted separately and the lines follow the mean of these replicates. Samples were taken at 0.583 days. Graph A, Dehydroascorbate reductase *(DHAR)* genes (squares are *DHAR (AT1G19550),* circles are *DHAR1 (AT1G19570:* probably a pseudogene (Noshi *et al.,* 2016b)), upward triangles are *DHAR2 (AT1G75270)* and downward triangles are *DHAR3 (AT5G16710);* Graph B, Monodehydroascorbate reductase *(MDAR)* genes (squares are*MDAR1 (AT3G52880),* circles are*MDAR2 (AT5G03630),* upwards triangles are*MDAR3 (AT3G09940),* downward triangles are*MDAR4 (AT3G27820)* and diamonds are*MDHAR6 (AT1G63940));* Graph C, Glutathione reductase (GR) genes(squares are *GR1 (AT3G24170),* circles are *GR2 (AT3G54660)).* Statistical significances of changes are shown in Table S3.

#### Regulatory genes

A range of genes have been identified that have been proposed to regulate ascorbate concentration (reviewed in (Bulley and Laing, 2016a)). We examined the expression of these genes in response to growth PFD, time of day and step change in PFD as for other genes described above. Note that *AMR1 (Zhang et al., 2009) (AT1G65770)* was not expressed in any treatment in our experiment.

#### Response to growth PFD

Only genes that showed a significant response to growth PFD are plotted in Fig. 11. These are *KONJAC (Sawake et al., 2015) (AT1G74910), SlHZ24 (Hu et al., 2016) (AT3G01470)* and *Dof22 (Cai et al., 2016) (AT3G47500).* The other genes showed little response to growth PFD (Table S5) suggesting they do not regulate ascorbate concentration in response to PFD at the transcriptional level.

**Figure 11.**
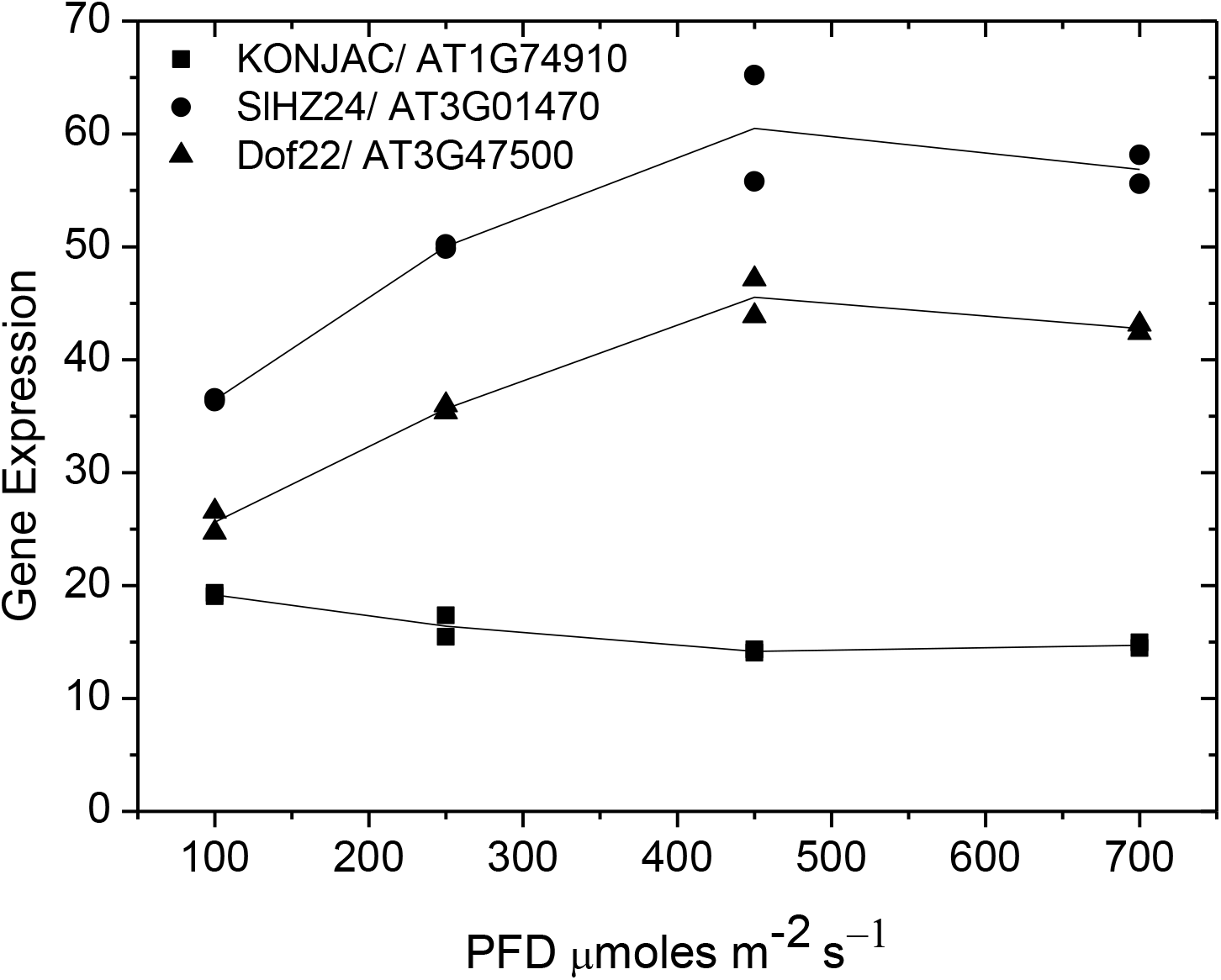
The response of gene expression of ascorbate regulatory genes to PFD. The two biological replicates are plotted separately and the lines follow the mean of these replicates. Square symbols represent *KONJAC (AT1G74910),* circles *SlHZ24 (AT3G01470)* and upward triangles *Dof22 (AT3G47500).* The data for all regulatory genes is shown in Table S6.

#### Response to time of day (diurnal trends)

Five genes showed significant responses to the time of day, three showing clear maxima during the night (Fig. 12A) and two maxima during the day (Fig. 12B). Other genes showed little change during the day/night cycle and are summarised in Table S6.

**Figure 12.**
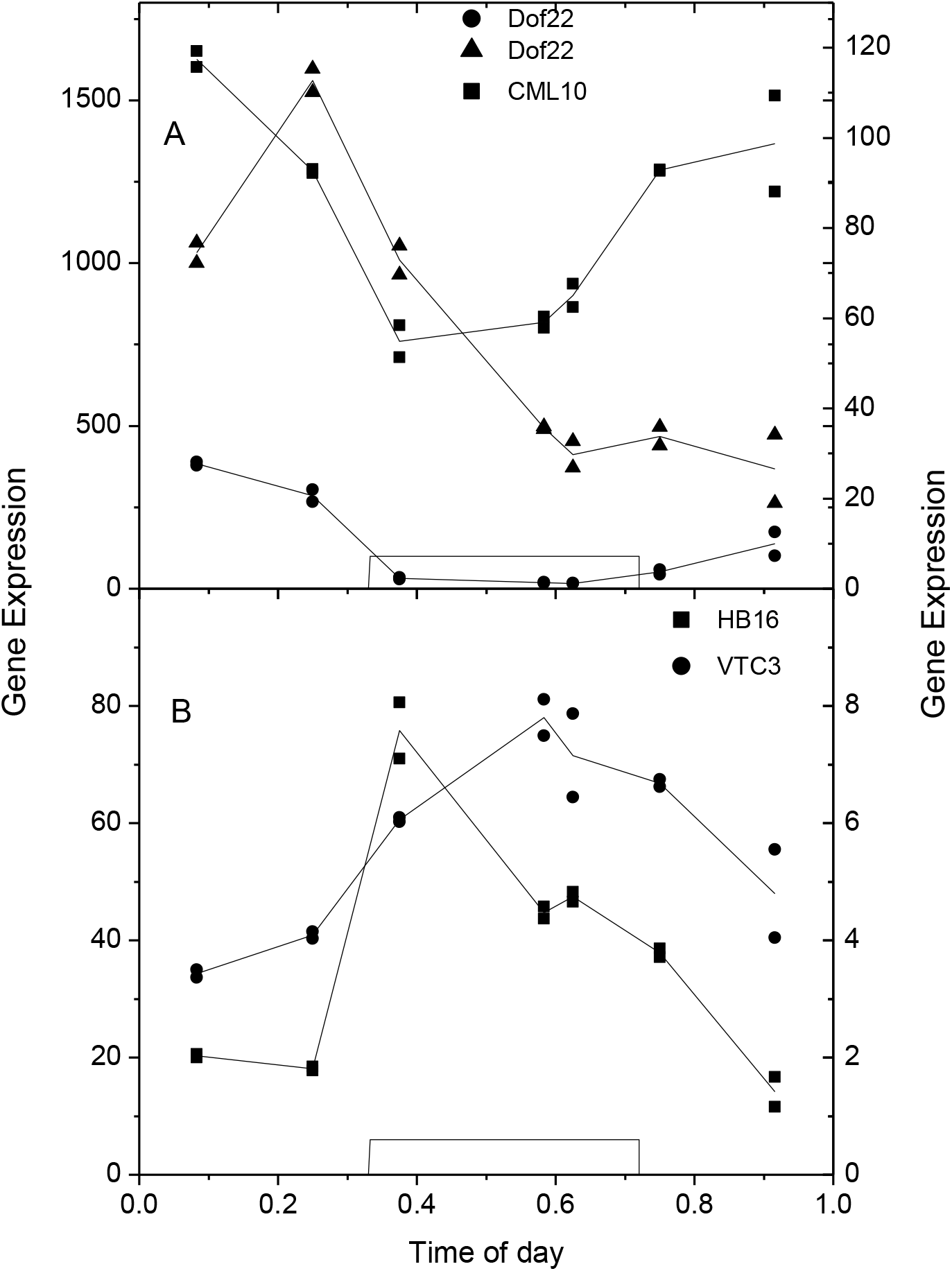
The response of gene expression of ascorbate regulatory genes to time of day. The two biological replicates are plotted separately and the lines follow the mean of these replicates. Plants were grown at 250 μmoles photons m^-2^ s^-1^: The square wave in graph A shows when the lights were on. Graph A. Genes showing a maximum during the dark period. Square symbols *CML10 (AT2G41090);* Circles *Dof22 (AT5G39660);* Triangles *Dof22 (AT3G47500)* (both right hand axis). Graph B Squares, *HB16 (AT4G40060);* Circles *VTC3 (AT2G40860)* (Right hand axis). Full data is shown in Table S7.

#### Response to a step change in PFD

Generally, regulatory genes showed a less than two fold change in gene expression upon increasing the PFD and in two cases the initial trend was a decrease in gene expression compared to the higher steady state expression of plants grown constantly at the higher PFD (Table S7). For example, *SlHZ24 (AT3G01470),* and *HB16 (Belmonte and Stasolla, 2009) (AT4G40060)* decreased in gene expression. Only *Dof22 (AT3G47500)* showed an upward trend towards the higher PFD value.

### Other ascorbate related genes

We selected a range of genes with ascorbate in their description from TAIR. These included ascorbate peroxidases, ascorbate oxidases, various transporters and others. While several of the genes, including ascorbate peroxidases, increased with increasing growth PFD, others decreased (see Fig. S8). Similarly, while some peroxidases showed a moderate (< 2 fold change) in diurnal gene expression, nothing outstanding was apparent. Interestingly on a change in PFD from 100 to 700 μmol m^-2^ s^-1^, several genes overshot the 700 μmol m^-2^ s^-1^ growth conditions and then showed some recovery.

## Discussion

This study focussed on aspects of the *Arabidopsis* growth environment that would perturb ascorbate biosynthesis and allow us to explore control of ascorbate biosynthesis at the transcriptional level. With this in mind, we grew *Arabidopsis* at PFDs from 100 to 700 μmol m^-2^ s^-1^, the higher levels being in the stressful photoinhibitory range (e.g. (Bailey *et al.,*2001)) especially at low night temperatures. These conditions were expected to promote high concentrations of ascorbate (e.g. (Dowdle *et al.,* 2007)), which we observed (Fig. 2A). Previous studies have reported that ascorbate shows a diurnal response, generally maximum around midday (Dowdle *et al.,* 2007; Massot *et al.,* 2012). This response was not observed (Fig. 2B)). Lastly, step increases in growth PFD have been shown to increase ascorbate (e.g. (Dowdle *et al.,* 2007)) which we did observe (Fig. 3B) but these were minimal compared to the literature (e.g. (Dowdle *et al.,* 2007). The differences between what we observed and the literature probably reflect differences in the details of the experimental procedures. Our plants were well expanded mature plants grown under short days to inhibit flowering. We validated and verified our RNAseq gene expression data by comparison to expected responses from the literature. We extracted RNA separately for the 4 to 6 replicate plants harvested for each treatment, and then combined the RNA to created two biological replicates. These replicates showed good agreement and allowed analysis of variance of the data, showing significant changes to occur. In order to further validate the results, we chose a range of genes with either published or expected responses to PFD and time of day. Our results confirmed published values (e.g. Fig. 4 and Fig. S1 and Fig. S2). We believe there is no further need for validation of this data. Note the ascorbate data is based on all 4 to 6 replicates being extracted and measured separately.

Ascorbate concentrations in leaves would be a function of biosynthesis (controlled by glucose supply and biosynthetic enzyme amount and activity), demand from side pathways (e.g. fucose and mannan biosynthesis from GDP-mannose, RG-II from GDP-galactose), by oxidation of ascorbate (ascorbate peroxidases and oxidases) and its recycling, by degradation, especially of the oxidised form (dehydroascorbate and monodehydroascorbate) and transport ofascorbate. This is illustrated in Scheme 1. Little is known about genes and enzymes that control transport (Maurino *et al.,* 2006) or degradation (Dewhirst and Fry, 2015), except in the chloroplast (Miyaji *et al.,* 2015) but the genes and enzymes involved in the L-galactose pathway of biosynthesis and the oxidation and recycling of ascorbate are well known. While we do not examine other potential pathways of ascorbate biosynthesis in this paper, as discussed earlier, the L-galactose pathway of ascorbate biosynthesis is the dominant or only pathway of ascorbate biosynthesis in *Arabidopsis.* The evidence for this is that the double GGP mutant (VTC2 and VTC5) is unable to grow past germination without L-galactose downstream substrate (Dowdle *et al.,* 2007). While it is possible that other pathways come into play in mature plants, this seems to us unlikely. In addition, the genes involved in these proposed alternative pathways are not all identified.

**Scheme 1.**
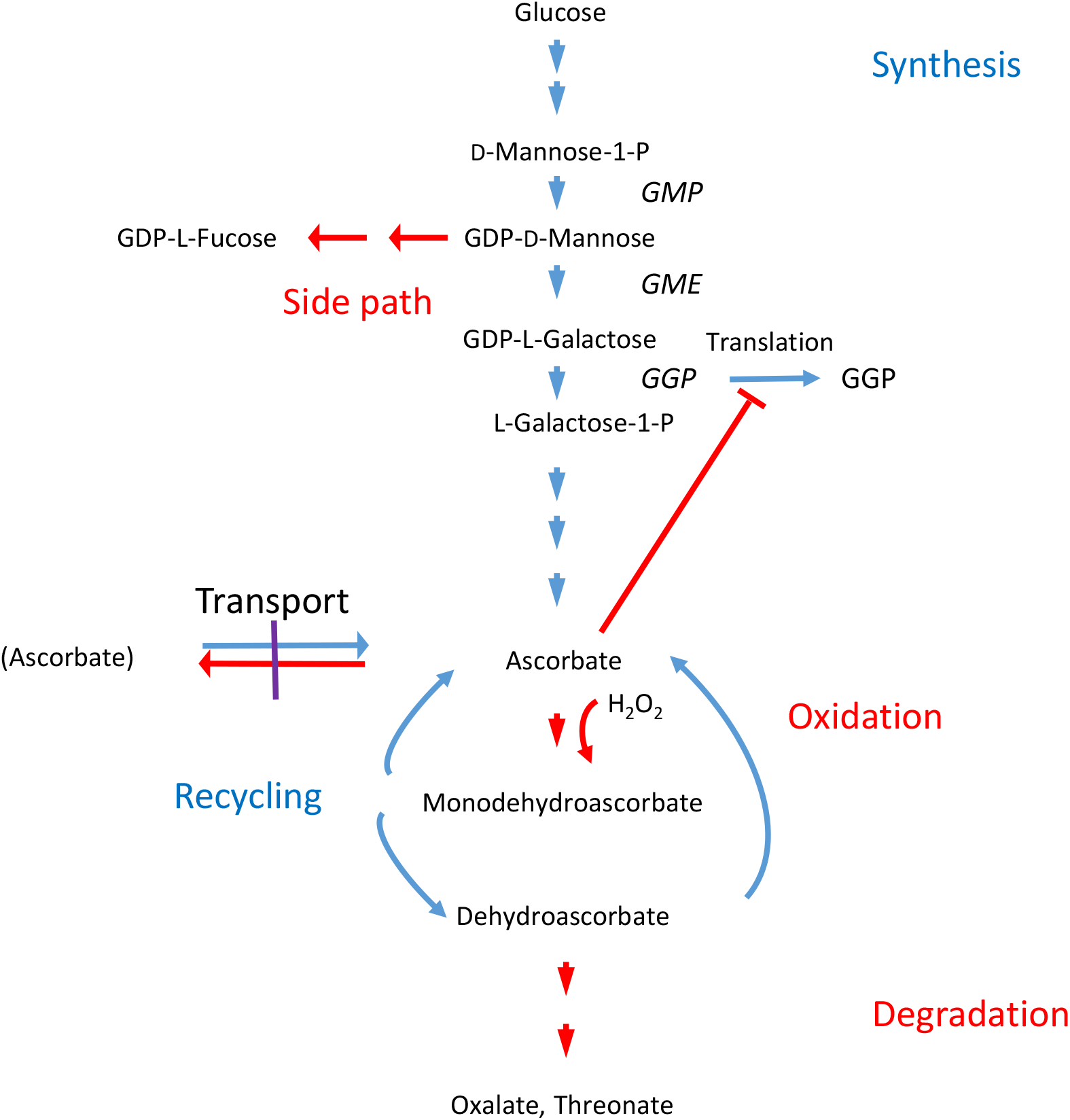
Outline of the various processes that determine ascorbate concentration. These include synthesis from glucose(including enzyme amounts and feedback inhibition of GGP translation by ascorbate)), competition from side paths(only one shown here), oxidation of ascorbate (symbolised by H2O2), recycling of oxidised ascorbate back to ascorbate and degradation of oxidised ascorbate to threonate and oxalate. Only key processes are shown. The verticalpurple line represents the cell membrane, the red line implies inhibitory, competitive or destructive processes,and the blue line synthetic processes. Factors that promote transcription, enzyme turnover (Bulley and Laing, 2016a) etc. are not shown.

Growth PFD and night temperature were the factors that most strongly influenced ascorbate concentration in our experiments, with higher ascorbate being found at lower night temperature. As the change from lights off to lights on occurred as a sharp transition, and the transition between the 5°C night and the 23°C day took 120 minutes (with lights on/off 60 minutes into this transition), the plants would have experienced 14°C day at both lights on (rising temperature) and lights off (falling temperature). Consequently, these plants would have been stressed with the combination of low temperature and high PFD. In spite of this, plant growth increased with growth PFD at both night temperatures (Fig. 1) although more at the higher night temperature. Ascorbate concentrations rose with increasing PFD (Table 1 and Fig. 2) reaching saturation at 450 μmol m^-2^ s^-1^. *GGP* also showed a strong response to PFD again saturating at 450 μmol m^-2^ s^-1^ (Fig. 5). Other genes in the L-galactose pathway either showed little or no response to PFD. Of the recycling genes, *DHAR1,* several *MDARs* and *GR1* all increased with increasing growth PFD (Fig. 8). Only three of the regulatory genes identified showed changes with growth PFD (Fig. 11). Interestingly, *KONJAC* which has been reported to stimulate GMP activity (Bulley and Laing, 2016a) decreased with increasing PFD, while *HZ24* (transcription factor enhancing biosynthesis genes (Bulley and Laing, 2016a)) and *Dof22* (negatively regulating ascorbate (Cai *et al.,* 2016)), both increased with increasing PFD even though they appear to have opposite effects. Expression of several ascorbate peroxidases *(APX)* increased with increasing PFD (data not shown).

We looked at correlations and relationships between ascorbate concentration and gene expression for plants grown at the four PFD levels. As the growth PFD increased, the concentration of ascorbate in leaves increased and we found several genes displayed good correlation between gene expression and ascorbate, especially early biosynthetic genes *(PMM, GMP, GME* and GGP) as well as some versions of *DHAR, MDAR* and *GR,* and several of the regulatory genes (Table S9). However, when the slope normalised ascorbate (i.e. ascorbate divided by the maximum ascorbate to give a scale of 0 to 1) was plotted against normalised gene expression, many fewer genes showed a close to 1:1 relationship. These included *GGP*, two *DHAR* genes and one *GR,* and no regulatory genes. This more stringent test for a simple functional control supports the idea that *GGP* is a central regulatory gene in ascorbate biosynthesis and that only some recycling genes are critical. The lack of a strong 1:1 relationship for regulatory genes suggests they may perform a role to fine tune ascorbate concentration but are not primary drivers of ascorbate concentration.

A second test we undertook was to sample plants over the diurnal cycle to look for evidence of ascorbate related genes being differentially controlled. Interestingly, ascorbate showed no diurnal change (Fig, 2B), with the 5°C night grown plants showing a consistently higher ascorbate than the 23°C night grown plants. This was in spite of their being a strong diurnal rhythm for *GGP*, and to a lesser extent *GMP* and *GME* (Fig. 6B). These genes all showed maximal gene expression before the light came on and then decreased later in the afternoon to prepare for the demands of ascorbate during the light period through generation of reactive oxygen species in photosynthesis. Studies in *Nicotiana benthamiana* using a hairpin to *GGP* showed that ascorbate dropped over several days when transcription of this key gene was reduced (Laing *et al.,* 2015). This suggests that the GGP protein must also turn over relatively fast and helps explain the strong diurnal rhythm in GGP expression. In contrast, *GDH* (Fig. 6B) and perhaps *GPP* (Fig. 6A), which also showed a muted diurnal trend, were maximal late in the light period (day). Of the recycling genes, only the GR genes showed any sign of a diurnal expression, again being minimal late in the day. The constancy of ascorbate over the day, in spite of the changes in gene expression, demonstrates the robustness of the overall control system to maintain ascorbate. Gene transcription is only one aspect of the regulatory system, translation, protein degradation and lifetimes of messenger RNA and proteins would also play a role.

The regulatory genes showed two patterns, either being minimal during the day (Fig. 12A) or maximal (Fig. 12B). Interesting, the negatively regulating *Dof22* genes were lowest when ascorbate demand was highest, whereas *CML10,* which promotes GMP activity, was also lower during the day. VTC3 showed a maximum during the day (Fig. 12B). This gene is suggested to function post-transcriptionally and to allow responses to heat and light (Conklin *et al.,* 2013). This is in agreement with its highest expression during the day.

The last experiment was to expose low light grown plants to high light. This has previously been shown to result in rapid and significant increases in ascorbate concentration (Dowdle *et al.,* 2007). However, we only observed a small increase after about 30 h (Fig. 3), although there was a long 14 h dark period in this time. This is in spite of the gene expression of *GGP* and to a lesser extent *GMP* increasing within this time. Of the recycling genes, *DHAR1* again increased strongly, and the *GRs* as well. In the case of regulatory genes, the regulatory genes showed a muted response, with little change after a step change in PFD (Table S7). We interpret the small increase in ascorbate after a large increase in PFD as reflecting the balance between synthesis and degradation. Thus the ability of the leaf to maintain and even increase ascorbate, with strong increases in *GGP* expression as well as recycling genes, shows both synthesis and recycling were increased but unable to exceed degradation within the short time span of this experiment. Obviously in the long term (2 weeks in this case) ascorbate concentrations come to a new PFD determined equilibrium.

In conclusion, we interpret our results to show that genes for *GGP* and to a lesser extent *GME* and *GMP* are critical in the control of ascorbate biosynthesis, and the recycling of ascorbate is important to maintain it in the reduced state. The muted and sometimes contradictory results with regulatory genes shows that the regulation of ascorbate biosynthesis by factors other than biosynthesis and recycling genes is still not clear.

## Acknowledgements

We would like to thank Jonathan Crawford for growing and maintaining the plants and Peter McAtee for advice on analysis of the transcript data using Galaxy

